# Synaptotagmin-7 participates in the regulation of acetylcholine release and short-term presynaptic facilitation in splanchnic nerve terminals

**DOI:** 10.1101/2020.04.13.039768

**Authors:** René N. Caballero-Florán, Mounir Bendahmane, Julie M. Philippe, Alina Morales, Arun Anantharam, Paul M. Jenkins

## Abstract

Disturbances that threaten homeostasis elicit activation of the sympathetic nervous system (SNS) and the adrenal medulla. The effectors discharge as a unit to drive global and immediate changes in whole-body physiology. Descending sympathetic information is conveyed to the adrenal medulla via preganglionic splanchnic fibers. These fibers pass into the gland and synapse onto chromaffin cells, which synthesize, store, and secrete catecholamines and vasoactive peptides. While the importance of the sympatho-adrenal branch of the autonomic nervous system has been appreciated for many decades, the mechanisms underlying transmission between presynaptic splanchnic neurons and postsynaptic chromaffin cells have remained obscure. In contrast to chromaffin cells, which have enjoyed sustained attention as a model system for exocytosis, even the Ca^2+^ sensors that are expressed within splanchnic terminals have not yet been identified. This study shows that a ubiquitous Ca^2+^-binding protein, synaptotagmin-7 (Syt7), is expressed within the fibers that innervate the adrenal medulla, and that its absence can alter synaptic transmission in the preganglionic terminals of chromaffin cells. The prevailing impact in presynapses that lack Syt7 is a decrease in synaptic strength and neuronal short-term plasticity. Evoked excitatory postsynaptic currents (EPSCs) in Syt7 KO preganglionic terminals are smaller in amplitude than in wild-type synapses stimulated in an identical manner. Splanchnic inputs also display robust short-term presynaptic facilitation, which is compromised in the absence of Syt7. These data reveal, for the first time, a role for any synaptotagmin at the splanchnic-chromaffin cell synapse. They also suggest that Syt7 has actions at synaptic terminals that are conserved across central and peripheral branches of the nervous system.

## Introduction

The term “fight-or-flight” refers to the state of heightened physiological and mental arousal triggered by a physical threat, emotionally charged event, or metabolic disturbance. Although the precise reaction to each stressor may vary, all share some basic characteristics of sympathetic activation. The adrenal medulla is a core effector of the sympathetic nervous system in the periphery [8]. When activated, it discharges a cocktail of powerful hormones (epinephrine, norepinephrine, and vasoactive peptides) into the suprarenal vein for circulation throughout the body [54]. These hormones modulate cardiac, pulmonary, and metabolic functions in ways that favor survival or preserve internal conditions when they are likely to be disturbed [8, 19, 20]. Thus, many of the “emergency” measures encompassed in the term fight-or-flight, rely on the adrenal medulla.

Secretion from the adrenal medulla is dependent on input from preganglionic, sympathetic fibers which pass into the gland via the splanchnic nerves [15, 21]. The fibers terminate on adrenomedullary chromaffin cells which secrete hormones contained in dense core granules [9]. Due to their experimental accessibility, a great deal is now known about the mechanisms underlying dense core granule exocytosis in chromaffin cells [1, 3]. On the other hand, little is known about the molecular operation of exocytosis in splanchnic neurons.

The purpose of this study was to characterize the role of one synaptic protein in particular – synaptotagmin-7 (Syt7) – in regulating exocytosis at the splanchnic-chromaffin cell synapse. Syt7 belongs to a large family of proteins, numbering 17 in total, many of which couple calcium influx to vesicle fusion [41, 45, 48]. Our interest in Syt7 at splanchnic terminals has developed as a result of recent discoveries concerning its function at central nervous system (CNS) synapses. There, Syt7 is now generally acknowledged to regulate synaptic transmission in important ways, and to be required for forms of synaptic plasticity that are driven by subtle variations in calcium levels, including asynchronous release and facilitation [2, 30, 36, 38, 39, 49]. To date, no synaptotagmins have been identified in splanchnic neurons.

The experiments described here were performed on adrenal slices obtained from wild-type (WT) animals or animals in which the *Syt7* gene had been deleted (hereafter referred to as Syt7 KO) [11]. The absence of Syt7 had several readily identifiable effects on splanchnic synaptic transmission, which was evoked by stimulating preganglionic input to chromaffin cells via electrical stimulation. Specifically, evoked EPSCs in KO slices are smaller in amplitude than those in WT slices. Decreases in the coefficient of variation (CV^-2^) in evoked EPSCs in KO slices compared to WT, without changes in the decay, is consistent with a presynaptic modulation of acetylcholine release probability that involves Syt7. Moreover, facilitation, which is ordinarily a robust property of this synapse, was abrogated in the absence of Syt7. The magnitude of tonic or basal currents on which synchronous EPSCs are imposed was substantially smaller in KO synapses than in WT synapses. These functional data are supported by fluorescent imaging of adrenal sections, in which Syt7-positive puncta are found to be associated with a marker of splanchnic neurons, choline acetyltransferase (ChAT).

In sum, we demonstrate here that Syt7 is indeed expressed within the splanchnic fibers that innervate the adrenal medulla and has a role in evoked release and in facilitation at their synaptic terminals. These data are the first to document a function for Syt7 in regulating short-term synaptic plasticity in the autonomic nervous system.

## Materials and methods

### Animals

Litters of adult male and female *Syt7*^-/-^ (gift of Dr. Joel Swanson; [11]) and *Syt7*^*+/+*^ (from a C57BL/6J background and obtained from Jackson Labs, Bar Harbor, ME) were used in these studies. Animals were group housed (2 to 5 per ventilated cage) with 24 hr (12/12 dark/light cycle) access to food and water. All animal procedures and experiments were conducted in accordance with the Institutional Animal Care and Use Committee. No randomization was performed to allocate subjects in the study.

### *In Situ* Electrophysiology Recordings and Analysis

*Syt7* +/- mice were crossed to generate *Syt7* -/- or +/+ littermates. Mice were genotyped according to instructions provided by Jackson Labs (https://www.jax.org/strain/004950). All electrophysiological studies were performed on littermates by experimenters blinded to the genotype of the animal.

3-4-month-old animals (male and female; 18 to 25 g) were gas anesthetized using an isoflurane drop jar technique and sacrificed by guillotine decapitation (all procedures are in accordance with approved UM IACAC protocol PRO000009265). Chromaffin cells are responsible for releasing catecholamines in response to stress, including hypoxia. Hence, isoflurane is used to induce a faster loss of consciousness compared to CO_2_ euthanasia (30 s to 1 minute versus several minutes) and reduce animal stress.

Adrenal glands were then quickly removed from the kidney and placed in ice cold (4°C) slicing solution containing, in mM: 62.5 NaCl, 2.5 KCl, 1.25 KH_2_PO_4_, 26 NaHCO_3_, 5 MgCl_2_, 0.5 CaCl_2_, 20 Glucose and 100 Sucrose (pH maintained at 7.4 by saturation with O_2_/CO_2_, 95/5% respectively) at an osmolarity of ∼ 315 milliosmolar (mOsm). Glands were subsequently embedded in 3.5% agarose block solution at 4°C. Approximately 300 μm thick sections were cut with a microtome (VF-300, CompresstomeTM; Precisionary instruments, Natick MA). Slices were transferred to a stabilization chamber where they were maintained at room temperature for 60 min in artificial cerebrospinal fluid (ACSF) containing in mM: 125 NaCl, 2.5 KCl, 1.25 KH_2_PO_4_, 26 NaHCO_3_, 1 MgCl_2_, 2 CaCl_2_ and 20 Glucose, pH 7.4 (with 95% O_2_ and 5% CO_2_ bubbling through the solution, ∼300 mOsm). Then individual slices were transferred to the microscope to the recording chamber (∼300 μL volume) continuously super-fused with ACSF (1-2 mL/min) at room temperature.

The adrenal gland was visualized at in the microscope (Nikon Eclipse FN-1) at X10 to determine the recording and stimulation areas. The cholinergic nerve terminals of preganglionic neurons were activated using a focal stimulating FHC tungsten metal electrode (2-3 MΩ). The electrode was placed in the border between the adrenal cortex and the adrenal medulla around 50 - 100 μm away from the recording pipette.

Recording micropipettes were pulled (P-97; Sutter Instruments, Novato, CA) from borosilicate glass capillaries (1.5 mm O.D.; Harvard Apparatus, Holliston, MA) for a final resistance of 3-6 MΩ. Pipettes were filled with a cesium-based internal solution of the composition in mM: 135 CsCl, 4 NaCl, 0.4 GTP, 2 Mg-ATP,0.5 CaCl_2_, 5 EGTA and 10 HEPES pH 7.25 – 7.3 (290 mOsm). Currents were recorded with Axon Instruments, Multiclamp 700B (Axon Instruments, Union City, CA), low pass filtered at 2 kHz. Chromaffin cells in medullary slices were identified using a Nikon Eclipse FN-1 microscope with a X40 water-immersion objective and a DAGE-MTI IR-1000 video camera. Whole-cell recordings (more than 8 GΩ before break-in) were obtained in voltage-clamp configuration, acquired at 2 kHz fixing the voltage at -20 mV. Series resistance was monitored throughout the experiment and experiments were aborted if changes greater than 20% occurred. The cells were chosen according to the access resistance and visual examination of their membranes. The EPSCs were evoked by stimulating the preganglionic input at 0.1 Hz and were distinguished by their all-or-none response to presynaptic stimulation and fast kinetics [4, 25, 52]. The reciprocal of the squared coefficient of variation (CV) of the synaptic response amplitude was quantified as (CV)^-2^ = 1/ [(SD/mean)^2^]. Disparities in this value is often interpreted as a change in quantal content due to a presynaptically mediated change in transmitter release [12, 34, 52]. Once a “synapse” was identified, to evaluate short-term synaptic plasticity the preganglionic input was stimulated at 5 – 50 V intensity (0.5 ms pulse duration) every 15 seconds at intervals of 60, 100, 200 and 500 ms or during high frequency trains (20Hz). In some experiments the calcium concentration in the ACSF was reduced 2mM to 0.5 mM, to avoid calcium saturation. Stimulation waveforms were introduced via a Grass S48 stimulator (Quincy, MA) that was triggered using Clampex9 software. Paired-pulse ratio (PPR) of EPSCs were calculated by dividing the amplitude of the second EPSC2 by that of the first EPSC1 (PPR = EPSC2/EPSC1); Differences assessed by PPR parameter will indicate changes in neurotransmitters release mediated presynaptically [7, 33, 55]. For experiments involving the application of hexamethonium (hexane-1,6-bis (trimethylammonium bromide)) a non-depolarizing nicotinic acetylcholine receptor (nAChR) antagonist at concentration of 100μM was added to the bath, and stimulation was conducted at 0.1 Hz (unless otherwise indicated) the slice was perfused with ACSF (1-2 mL/min) in presence of the drug for 5 minutes before washout and for before 5 min to obtain a baseline response. Peak current amplitudes were searched identified manually using pCLAMP 10 (Molecular Devices, San Jose, CA), and visually monitored to exclude the erroneous noise. The current response was fit by a single exponential equation to obtain the time constant (tau) of decay. Basal current (during high frequency stimulation) is measured as the difference between the sustained currents reached during the train and the overall baseline current of the record [39].

### Immunofluorescence

3-4-month-old mice (male and female) were administered a ketamine/xylazine mixture (80 mg/kg body weight ketamine and 10 mg/kg xylazine) via intraperitoneal injection. The mice were perfused by cardiac perfusion of PBS followed by 4% paraformaldehyde (PFA). Adrenal glands were immediately removed and fat and connective tissues surrounding the glands were trimmed off. Adrenal tissue was subsequently processed using a standard single-day paraffin preparation protocol (PBS wash followed by dehydrations through 70%, 95%, and 100% ethanol with final incubations in xylene and hot paraffin under vacuum) using a Leica ASP 300 paraffin tissue processor. Paraffin sections were cut 7 μm thick using a Leica RM2155 microtome and placed on glass slides. Tissue sections were deparaffinized and rehydrated using a standard protocol of washes: 3 × 4-min xylene washes, 2 × 2-min 100% ethanol washes, 2 × 2-min 95% ethanol washes, and 1 × 2-min 70% ethanol wash, followed by at least 5 min in ddH_2_O. Antigen retrieval was conducted for brain sections by microwaving the deparaffinized brain sections for 20 min in 10 μM sodium citrate in water. Sections were cooled, washed for 15 min in ddH_2_O rinsed in PBS for 5 mins, and blocked using blocking buffer (5% BSA, 0.1% Tween 20 in TBS) for 1 hour at room temperature. No antigen retrieval was performed for deparaffinized adrenal gland sections. Immediately following deparaffinization, adrenal glands were incubated with blocking buffer for 1 hour at room temperature. All slides were incubated overnight at 4°C with primary antibodies diluted in blocking buffer (Rabbit anti Syt7, Mouse anti TH, Goat anti ChAT, diluted 1:400). On the following day, slices were washed 3x for 15 minutes each with PBS containing 0.2% Tween 20 and incubated with secondary antibodies diluted in blocking buffer for 1hr at room temperature. Fluorescently conjugated secondary antibodies Alexa 488, 568, and 647 (1:250, Life Technologies) were used. Secondary antibodies were washed three times with PBS containing 0.2% Tween 20, for 15 minutes each and mounted on glass slides with Prolong Gold.

Imaging was performed using a confocal microscope (LSM880, Zeiss, Germany) with a 63x oil immersion objective in the sub-diffraction, Airyscan mode. Excitation was accomplished using 405-, 488-, 561-, and 633-nm lasers. All images were further processed in Adobe Photoshop CS6 software.

### Statistical Analysis

All data were analyzed using GraphPad Prism (Version 8.0) Software, San Diego CA, USA. The Shapiro-Wilk test was used to ensure normal (Gaussian) distribution of the samples followed by a Bartlett’s test to check the homogeneity of the variances based on the means. Changes in amplitude of EPSC and PPR were analyzed using two-way ANOVA followed by Sidak’s multiple comparisons test to compare differences between experimental conditions, or one-way ANOVA following by Dunn’s multiple comparisons test, as appropriate. When only two conditions were compared Student’s t-test or Mann-Whitney test was used. A significance level of 0.05 was used for all statistical tests. For box and whisker plots, boxes represent the 25-75% confidence interval, horizontal lines are the median value, the plus symbol represents the mean, and the whiskers show the full data range.

## Results

### Synaptotagmin-7 regulates exocytosis at the splanchnic-chromaffin cell synapse

To confirm the contribution of Syt7 in splanchnic-chromaffin cell synapse and determine whether a presynaptic mechanism was involved, we measured cholinergic synaptic strength in adrenal gland slices. A stimulation electrode was placed at the border between the cortex and medulla in an adrenal section prepared as described in the Methods (Figure 1A). Electrical pulses were applied to stimulate splanchnic processes while recording from chromaffin cells, which were voltage-clamped in the whole-cell configuration. Excitatory postsynaptic currents (EPSCs), evoked by splanchnic stimulation, were measured in glands obtained from both WT and Syt7 KO animals (Figure 1). These currents are reversibly inhibited by the nicotinic receptor antagonist, hexamethonium (Figure 1B) [52]. The amplitude of evoked EPSCs and charge transferred were significantly smaller in slices that lack Syt7 compared to WT slices (Figure 1C-E); However, no difference was detected in single exponential decay time constant of the EPSC (Figure 1F), indicating that the currents are mediated by a similar pool of acetylcholine receptors [4, 27]. Since single exponential fits of EPSC decay may not reflect changes in asynchronous release, we estimated the skewness of the data by normalizing charge transfer to peak EPSC amplitude (Figure 1G). We did not detect differences between WT and Syt7 KO, consistent with no change in asynchronous release. Deletion of Syt7 caused a significant decrease in the inverse of the coefficient of variation (CV^-2^) of synaptic current amplitude (Figure 1F) consistent with a presynaptic modulation of neurotransmitter release probability [12, 34, 52].

**Figure 1.**
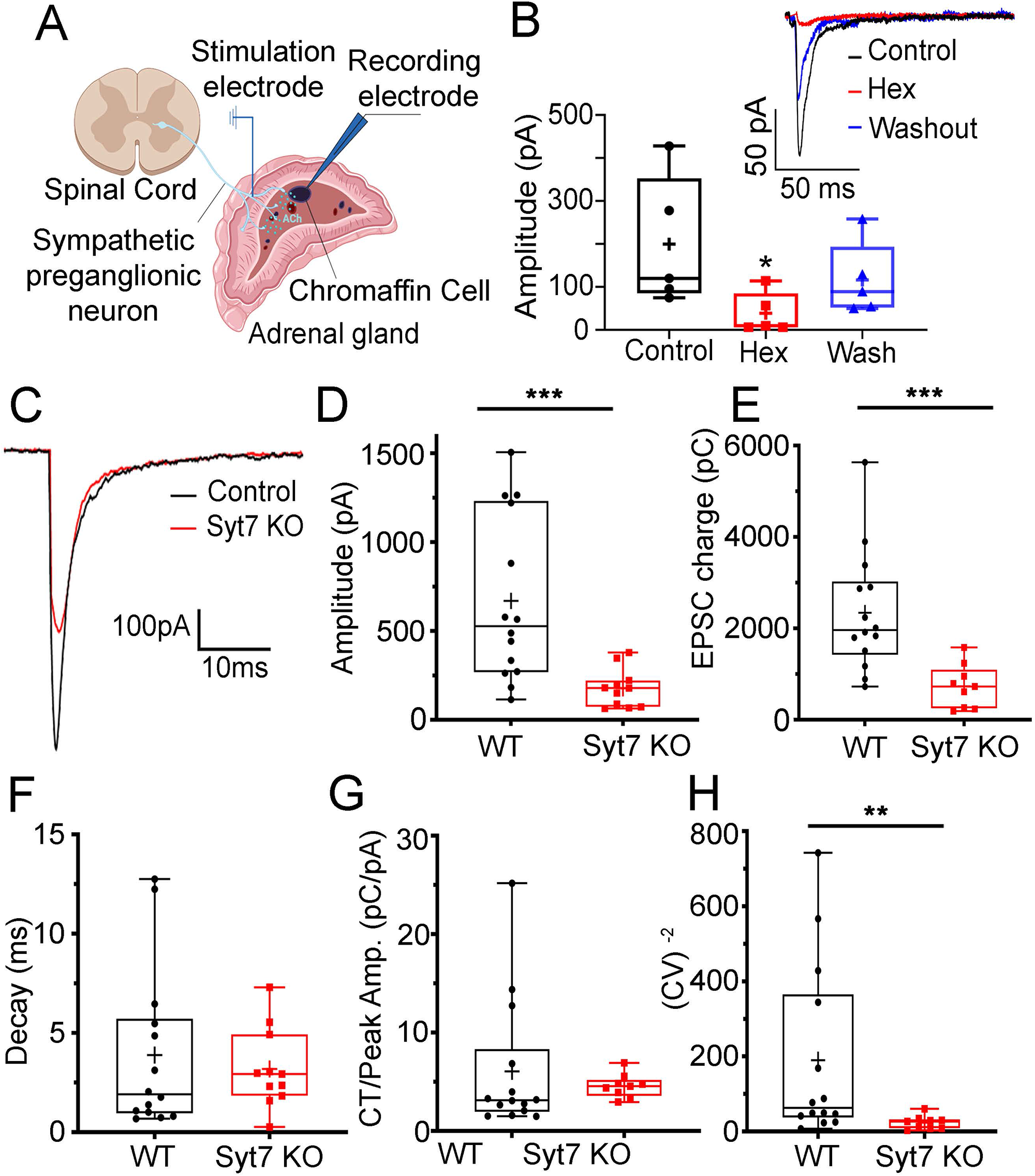
Comparison of evoked EPSCs in WT and Syt7 KO synapses. EPSCs were evoked by stimulating preganglionic input to the adrenal medulla with a bipolar stimulating electrode. Cartoons created with BioRender (www.biorender.com). **B**. EPSCs recorded in a chromaffin cell evoked by stimulating the preganglionic nerve terminals (black) were blocked by the cholinergic antagonist hexamethonium (red), and recovered during washout (blue), \**p*=0.032, Kruskal-Wallis, Dunn’s multiple comparisons (n=5 slices; 3 independent preps). Representative EPSCs at WT synapses: Control (black), during block by Hexamethonium (Hex, red), and after Washout (blue) (*inset*). **C**. EPSCs recorded in chromaffin cells by stimulating the WT (black) and Syt7 KO (red) preganglionic nerve terminals. **D**. Averaged peak amplitudes of evoked EPSCs from Syt7 KO (red) are decreased compared to WT (black). \*\*\**p*< 0.001, Mann Whitney test (n=14 WT and n=9 KO slices; > 6 independent preps). **E**. Average EPSC charge transfer from Syt7 KO (red) mice are decreased compared to WT (black) mice. \*\*\**p*< 0.001, Mann Whitney test (n=14 WT and n=9 KO slices; > 6 independent preps).**F**. Average decay time constants of evoked EPSCs in chromaffin cells after stimulation of WT (black) and Syt7 KO (red) axons. p=ns, Mann Whitney test (n=14 WT and n=9 KO slices; >6 independent preps). **G**. Charge transfer normalized to peak EPSC amplitude from evoked EPSCs are not different in WT (black) mice compared to Syt7 KO (red) mice. p=ns, Mann Whitney test (n=14 WT and n=9 KO slices; >6 independent preps) .**H**. CV^-2^ of the EPSC amplitude was significantly greater in chromaffin cells from WT (black) compared to Syt7 KO (red) mice. \*\**p*< 0.05, Mann Whitney test (n=14 WT and n=9 KO slices; > 6 independent preps).

### Paired-pulse facilitation is eliminated in synapses lacking Syt7

Syt7 has been reported to function as a specialized calcium sensor that mediates synaptic facilitation in several types of synapses in the brain [30, 50, 51]. The main purpose of this study was to test whether it functions in a similar capacity in splanchnic nerve terminals. Figure 2 shows that facilitation is indeed a property of synapses within the adrenal medulla. This was demonstrated by applying two successive depolarizing pulses and calculating the paired-pulse ratio (PPR, the amplitude of the second EPSC divided by the amplitude of the first). Interstimulus intervals (ISIs) ranging from 60 ms to 200 ms consistently resulted in PPRs above 1 (Figure 2A). On the other hand, PPRs above 1 were not observed at synapses lacking Syt7, irrespective of the ISI (Figure 2A). To rule out that this was not a consequence of the first pulse releasing so much transmitter that the terminals were already partly exhausted, we reduced evoked release by lowering extracellular calcium from 2.0 mM to 0.5 mM [30] (Figure 2B). Even under these conditions of low release probability, Syt7-deficient synapses did not exhibit facilitation.

**Figure 2.**
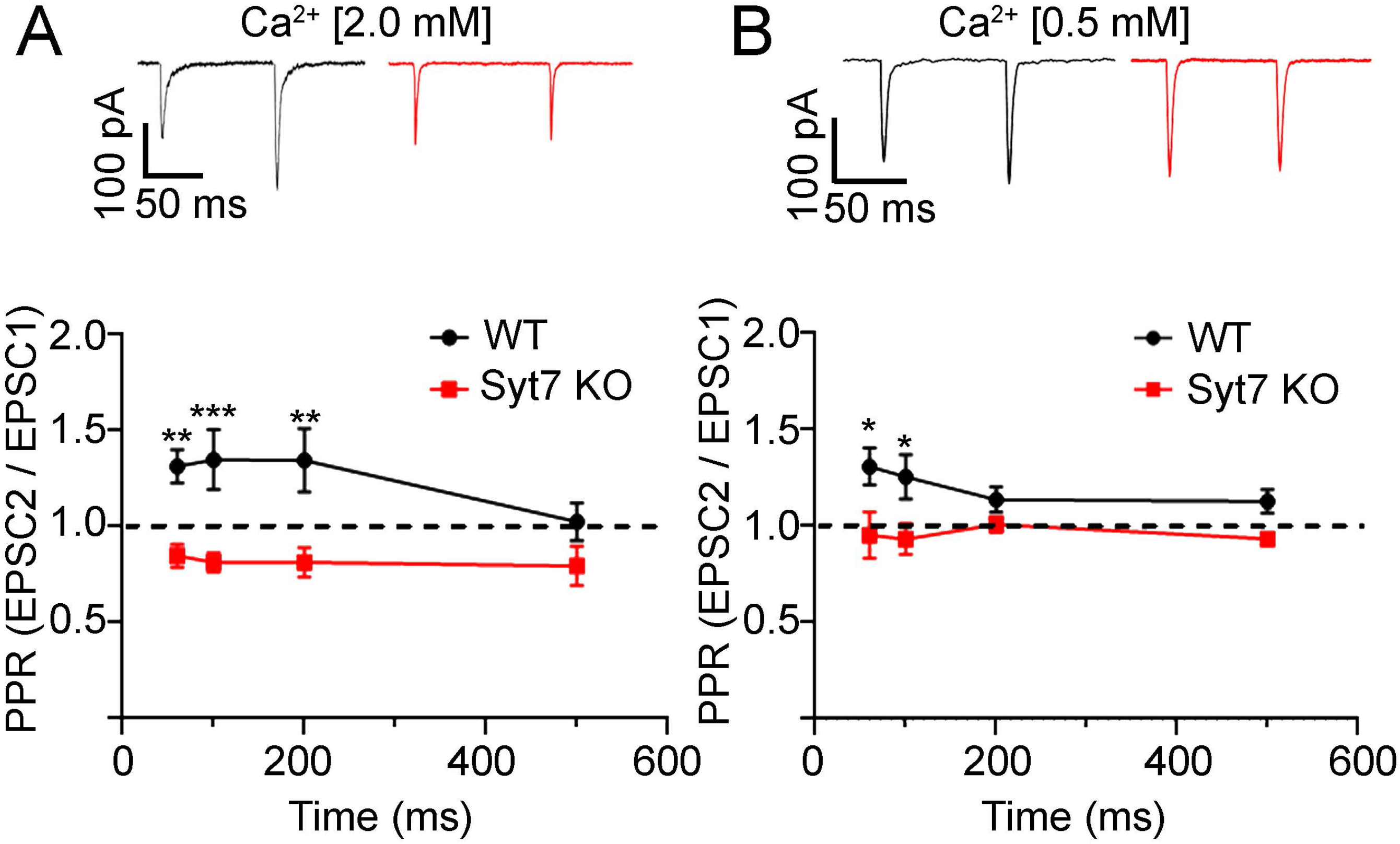
Synaptic facilitation is evident in WT but not Syt7 KO medullae. **A**. Representative traces (top) and averaged paired-pulse ratio (PPRs) ± SEM from evoked EPSCs at different interstimulus intervals (ISIs). PPRs at WT (black) and Syt7 KO (red) synapses are significantly different at ISIs of 60, 100 and 200 ms intervals but not at 500 ms. F (1, 79) = 35.76 \*\**p*=0.002, \*\*\**p*<0.001, Two-way ANOVA (n=13 WT and n=13 KO slices; >6 independent preps). **B**. Experiments were repeated at low extracellular Ca^2+^ (0.5 mM) and PPRs calculated as in A. \**p*=0.01, Two-way ANOVA (n=13 WT and n=15 KO slices; >6 independent preps).

### High frequency trains result in facilitating EPSCs in WT but not Syt7 KO synapses

Chromaffin cells of the adrenal medulla vary dramatically in the rate at which they fire [14, 22]. The variation in firing rate, in turn, may reflect the changing demands placed on them as effectors of the sympathetic stress response [18]. To model “full activation” of the medulla, a 20 Hz stimulus train was applied to WT and Syt7 KO splanchnic fibers [14]. Under these experimental conditions, two components are distinguishable: the synchronous release or phasic transmission, which is the first synaptic response riding on top of the plateau current, and the asynchronous component, also referred to as tonic transmission or basal current [2, 36-38, 42, 43]. Cholinergic release increased monotonically during the stimulus train (Figure 3A, C). The net increase in EPSC size may arise from several factors, including facilitation, receptor desensitization, and spillover [10]. Conversely, high frequency stimulus trains applied to Syt7 KO fibers result in a net decrease in EPSC size in chromaffin cells (Figures 3B, C, and E). Another notable difference between the records shown in Figures 3A and 3B is that the basal or tonic current, on top of which synchronous EPSCs ride, is markedly smaller in the absence of Syt7 (Figure 3D). The interpretation of the basal current is not simple and will undoubtedly require further research to clarify its mechanism in this terminal. However, this phenomenon has previously been attributed to an asynchronous neurotransmitter release component related to the increased likelihood of vesicle fusion, which can be compromised if Syt7 is absent [37].

**Figure 3.**
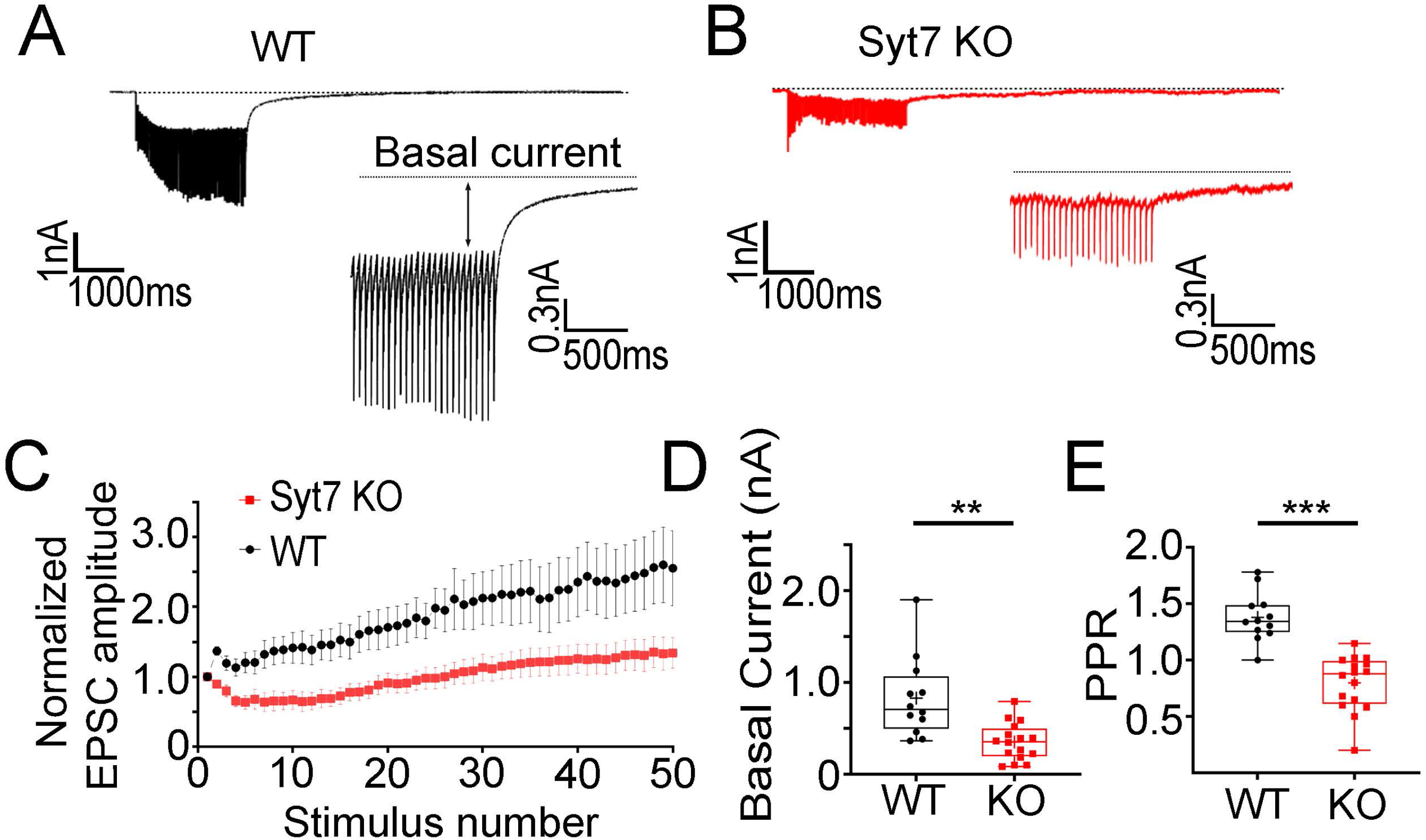
Basal current is reduced at Syt7 KO synapses. A, B. Synaptic responses to 20 Hz stimulation recorded from WT (black) and Syt7 KO (red) preparations. Expanded traces show the basal current. C. Summary graph of individual EPSC amplitudes normalized to the first EPSC amplitude during a train (WT in black and Syt7 KO in red). D. Summary graph of the average basal current, basal current amplitudes was significantly greater in chromaffin cells from WT (black) compared to Syt7 KO (red) mice\*\*\**p*<0.001, t=3.840, df=26 Student’s t-test (n=12 WT and n=16 KO slices; > 6 independent preps). (n=7 WT and n=11 KO slices; >4 independent preps). E. Average of PPRs calculated by dividing the second EPSC in the train by the first is. Facilitation is reduced in in chromaffin cells from Syt7 KO (red) conmpared with WT(black) mice \*\*\**p*=0.005, t=6.515, df=26 Student’s t-test (n=12 WT and n=16 KO slices; >6 independent preps

### Synaptotagmin-7 is present in preganglionic, cholinergic axons that innervate the adrenal medulla

Aspects of synaptic transmission in the adrenal medulla are compromised in animals lacking Syt7. Therefore, it was important to verify that Syt7 is localized to splanchnic terminals responsible for releasing ACh onto chromaffin cells. Immunocytochemical analysis was performed on adrenal sections which were exposed to antibodies for choline acetyltransferase (ChAT) and tyrosine hydroxylase (TH) – to label ACh-producing neurons and chromaffin cells, respectively – in addition to Syt7. ChAT-positive fibers and “varicosities” containing a cluster of fluorescent ChAT puncta, were frequently observed. The punctate appearance of ChAT immunofluorescence suggests it may be frequently associated with synaptic vesicles in splanchnic processes and boutons [17, 46, 47] In WT sections, ChAT clusters were sometimes enriched in Syt7. One such cluster is boxed in yellow in Figure 4A and expanded in Figure 4B. Syt7 immunofluorescence was barely detectable in adrenal medullae harvested from Syt7 KO mice compared to WT littermates (Figure 4A and B). Fluorescent puncta identified by the Syt7 antibody in KO medullae likely result from non-specific labelling of other intracellular material. This supposition is reinforced by Figure 4C, in which Syt7 immunoreactivity is absent from KO adrenomedullary lysates, as well as Figure 4D, which shows an almost total absence of fluorescence in Syt7 KO hippocampal sections probed with an anti-Syt7 antibody.

**Figure 4.**
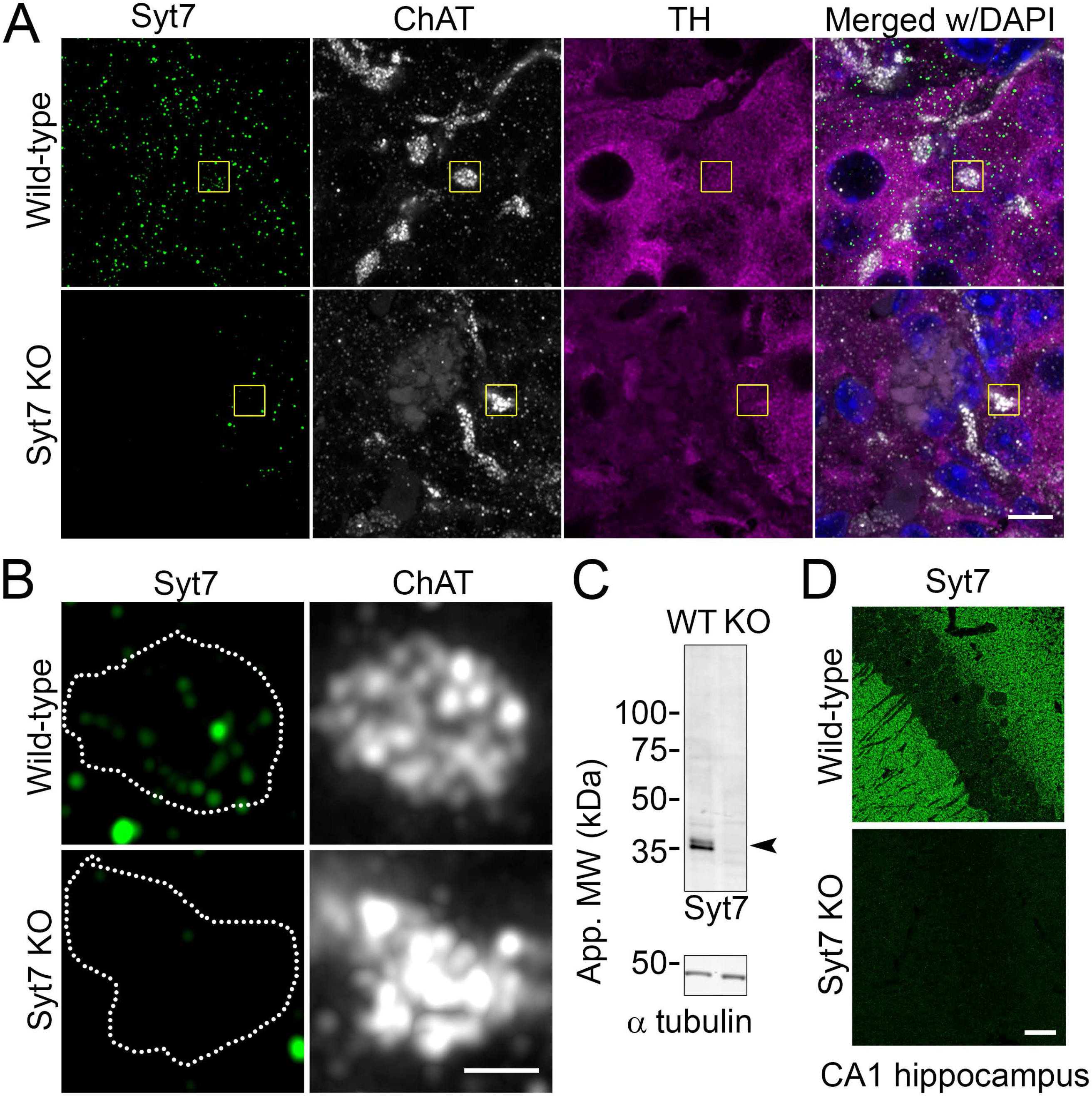
Syt7 is expressed in splanchnic processes and chromaffin cells in the adrenal medulla. **A**. Adrenal slices were immunostained with antibodies to choline acetyltransferase (ChAT) (white), tyrosine hydroxylase (TH) (magenta), and Syt7 (green). DAPI is included as a nuclear stain. Syt7 is co-localized with areas of ChAT and TH fluorescence, indicating that it is expressed in both splanchnic processes, as well as chromaffin cells, respectively. Scale bar, 5 □m. Some non-specific fluorescence is apparent in Syt7 KO slices exposed to the Syt7 antibody. Clusters of ChAT puncta were frequently observed in both WT and Syt7 KO slices. Two such regions are boxed (yellow) and expanded in **B. B**. A ChAT cluster enriched in Syt7 in a WT slice. A similar ChAT cluster in a Syt7 KO slice is also shown. Scale bar, 1 □m. **C**. Lysates from WT and Syt7 KO adrenal glands (2 in each lane) were run on a gel, transferred to nitrocellulose, and probed for Syt7 (top). Immunoreactivity for Syt7 is absent in Syt7 KO adrenal lysates. Alpha tubulin (bottom) was used as a loading control. Data are representative of 3 separate experiments. **D**. Representative images of the CA1 subfield of the hippocampus taken from coronal brain sections of wild-type (top) or Syt7 KO (bottom) mice immunoassayed with antibodies against Syt7 (green). Scale bar, 20 μm.

## Discussion

We have shown here that a ubiquitous Ca^2+^-binding protein, Syt7, is expressed within the neurons that innervate the adrenal medulla. These data are the first to implicate a role for any synaptotagmin in regulating neurotransmission at the splanchnic-chromaffin cell synapse. As is the case in the CNS, the functions of Syt7 in the periphery are closely tied to a property that sets it apart from the other Ca^2+^-binding members of the synaptotagmin family – its exceptionally high affinity for Ca^2+^ [6, 28]. What situations might demand such a Ca^2+^ sensor? Two closely related forms of synaptic plasticity are thought to rely on submicromolar Ca^2+^ - asynchronous release and facilitation [31, 44]. While our experiments initially do not address the possibility of asynchronous release, facilitation is clearly a robust property of the splanchnic-chromaffin cell synapse (Figure 2). And, consistent with published studies in central synapses [30, 51], facilitation is prevented in the absence of Syt7, whether it is driven by a pair of closely-spaced depolarizing pulses, or a high frequency stimulus train (Figures 2 and 3).

While the single EPSCs properties we describe here are consistent with those previously reported in identical gland preparations [4, 27, 32, 52], the decrease in evoked EPSCs amplitude in Sy7KO synapses compared with WT(Figure 1), was not expected in single stimulus experiments. This differs from what has been previously shown with respect to the impact of Syt7 on baseline synaptic responses [30, 36]. We did not detect differences in the decay time constant of EPSCs evoked by a single stimulus (Figure 1F), suggesting that a similar group of receptors is being activated in WT and Syt7 KO mice. The high degree of overlap in time course in the decay times between EPSCs in Syt7 vs WT indicates that transmitter vesicles are released with a high degree of synchrony when synapses are stimulated, and the fact that the evoked currents are prevented by hexamethonium indicate that they are acetylcholine-dependent [32, 52]. This is consistent with other studies that have reported changes in amplitude of stimulated currents, without concomitant changes in kinetics, likely due to alterations in the amount of neurotransmitter coming from presynaptic terminals [42, 43].

There are a number of potential explanations for the difference in baseline EPSCs in Syt7 KO splanchnic terminals. It may be that fewer presynaptic axons are activated by the stimulation electrode in Syt7 KO adrenal sections, possibly resulting from a different arrangement of fibers entering the medulla. However, chromaffin cells in situ are stimulated simultaneously by splanchnic nerve fibers, which are more relevant than gap-junctions, indicating that stimulation is likely an all-or-none event [32]. Altered EPSC amplitudes in the KO could also be due, in theory, to a reduction in postsynaptic nicotinic receptor expression. However, It was previously shown that nicotinic currents in dissociated WT and Syt7 KO chromaffin cells are not discernibly different, rendering such a possibility unlikely [5]. Note as well that robust synchronous EPSCs are still evident in Syt7 KO medullae, which does suggest other “fast” synaptotagmins (e.g., Syt1, Syt2, etc.) are likely expressed in splanchnic neurons and collaborate with Syt7 to regulate exocytosis.

Neurotransmitter release evoked with only a single pulse in WT mice is almost completely synchronous, but seriously compromised in Syt1-deficient mice, which still have considerable residual current upon one pulse stimulation. This small amount of residual current at the synapses in Syt1 KO cells is primarily asynchronous release and occurs at both excitatory and inhibitory synapses [42]. Therefore, in WT mice, even the evocation of currents by a single pulse can present an asynchronous release fraction, so it is possible that the amount of remnant neurotransmitter (asynchronous release) released in Syt1 absence, is the same that we are losing in the absence of Syt7, and it is for this reason that we detected a decrease in presynaptically evoked EPSCs amplitude from Syt7 KO compared with WT (Figure1). A similar situation occurs in Syt7/Syt2-deficient calyx synapses where evoked synchronous neurotransmitter release, showed decrease in amplitude of EPSCs evoked by isolated action potentials [39].

Another hypothesis would be that current amplitude decrease in Syt7 KO vs. WT it could be related with the postsynaptic feedback after stimulation and Ach receptors activation. Neuropeptide Y (NPY) is co-stored and co-released with the catecholamine’s in chromaffin cells and is required to maintain preganglionic-chromaffin cell synaptic strength, an effect mediated by activation of adrenal presynaptic Y5 receptors. NPY-KO mice was decrease in single pulse evoked EPSCs amplitude in adrenal slices from fasted mice [52], like our Syt7 KO condition (Figure 1). It could be interpreted that the difference in amplitude of EPSCs in Syt7 KO with respect to WT is influenced by a decrease in NPY release and therefore a lower activation of presynaptic Y5 receptors; due to the already diminished strength of the cholinergic preganglionic-chromaffin cell synapse in the Syt7 KO. In addition, the difference in the coefficient of variation (CV^-2^) in the amplitude of the synaptic current in single pulse stimulation (Figure 1), is consistent with the idea that the decrease in synaptic strength is due to a mechanism presynaptic where the probability of transmitter release is decreased in the absence of Syt7.

No evidence was found in this study for delayed release of neurotransmitter that persists after the end of a single action potential in our preparations as neither spontaneous nor miniature EPSCs were detected in our recordings. Spontaneous electrical activity in the slices of the adrenal gland is not routinely observed, unless it is caused by excitation by high concentrations of potassium [32, 52]. Syt7 is not on its own required for clamping spontaneous mini release or for mediating fast calcium-triggered release, consistent with previous studies on other synapses [2, 30, 38, 53]. Previous studies in CNS neurons have shown that overexpressed Syt7 can increase mini release[2]. Therefore, the clamping function by Syt7 may not be apparent under physiological conditions in the splanchnic synapse because Syt7 may not be expressed at sufficiently high levels, especially within presynaptic terminals.

Differences between WT and Syt7 KO synapses were detected beyond the synchronous component of the evoked EPSC. The basal current, on top of which synchronous EPSCs ride, was also substantially reduced in synapses that lacked Syt7. Although basal current has frequently been attributed to an asynchronous component of release (i.e., where secretion is not time-locked to the arrival of an action potential; [39]), its origins at the splanchnic-chromaffin cell synapse are not immediately obvious. For example, the basal current may report on the stimulation of chromaffin cells by neurotransmitter that has “spilled over” from neighboring terminals – a phenomenon that has been extensively studied at central synapses [10, 16, 35] – or it may reflect the direct activation of non-canonical (e.g., GPCR-dependent), slow postsynaptic conductances known to be active in the medulla [25]. Splanchnic neurons are known to house and secrete a multitude of peptide cargos [22]. We cannot yet account for the various ways, subtle or otherwise, in which peptidergic neurotransmission contributes to the phenomena measured here. The basal current may also reflect electrotonic current spread from one depolarized chromaffin cell to another. Electric coupling is believed to be enhanced in conditions of increased splanchnic nerve activity [23, 26].

Although it is possible that desensitization of postsynaptic receptors could play a role in the decreases in basal current, previous studies have shown that desensitization does not play a major role in this synapse. at peripheral ganglionic synapses, acetylcholine receptor-mediated EPSCs can last up to ten minutes in the presence of neostigmine, an acetylcholinesterase inhibitor, without being desensitized, an effect that is blocked by DHBE, a nicotinic antagonist [13]. Similarly, recordings performed in the presence of a high concentration of potassium (25mM), also in adrenal gland slices, show that it is possible to measure EPSCs even with high neurotransmitter concentrations in the intrasynaptic space [32]. Both previous strategies likely involve the release of acetylcholine in much greater quantity than the stimulation of the short remaining fibers that we are stimulating in our preparation. We hypothesize that the changes in the phasic current as well as in the basal current, are both consequences of the decrease in the neurotransmitter amount coming from the presynaptic terminals where Syt7 is absent (Figures 3 and 4). Clearance of by transporters may play an important role in this process.

Overall, this study provides strong evidence that basic functions of splanchnic-chromaffin cell synapse operation depend on Syt7, including a form of short-term synaptic plasticity termed facilitation. It has been suggested that facilitating synapses signal high frequency information to target cells, thereby modulating their excitability [29]. However, the physiological role of facilitation has remained elusive. In the context of the sympatho-adrenal system, facilitation may have a role in amplifying epinephrine discharge from chromaffin cells during conditions that increase sympathetic tone, including hypoglycemia [40, 52]. The resulting increase in circulating epinephrine would then be expected to increase blood glucose via multiple metabolic pathways [24, 40, 52]. A hypothesis, which future studies should test, is that regulated physiological responses to metabolic stressors (e.g., fasting) will require release driven by splanchnic Syt7. Such studies may have to wait until Syt7 expression can be abrogated solely in the periphery, and in a tissue-specific manner. In fact, tissue-specific deletion of Syt7 will be necessary to definitively disentangle its functions in controlling CNS drive of the sympathetic nervous system, from pre- and post-synaptic functions of Syt7 in the adrenal medulla. While these sorts of efforts will not be trivial, the data presented here encourage deeper investigations into the molecular mechanisms of release at these and other autonomic synapses about which very little is known.

## Acknowledgements

We would like to thank Dr. Will Birdsong (University of Michigan) and Dr. Kevin Bender (UCSF) for critical discussion of our data.

